# Maternal and early-life longitudinal cytokine profiles in a South African birth cohort: the impact of HIV

**DOI:** 10.1101/2025.10.14.682293

**Authors:** Tusekile S. Kangwa, Catherine J. Wedderburn, Jenna F. Annandale, Francesca Little, Heather J. Zar, Dan J. Stein, Petrus J.W. Naudé

## Abstract

**Background:** During pregnancy, exposure to maternal HIV and a disrupted cytokine environment may impact foetal immune development and health outcomes through cytokine-mediated mechanisms. We evaluated (i) peripheral blood cytokine differences in pregnant women with and without HIV, (ii) longitudinal differences in HIV-exposed uninfected (HEU) and HIV-unexposed uninfected (HUU) children, and (iii) latent cytokine groupings. Additionally, we explored the impact of maternal antiretroviral treatment (ART) initiation timing on cytokine levels.

**Methods:** We assessed 399 mother-child pairs in the Drakenstein Child Health Study (DCHS), in pregnancy (n=179 mothers with HIV and n=220 without HIV) and their children at 6 weeks, 2-, 3-, and 5-years. Eighteen serum immune markers were quantified with ELISA and multiplex assays. Group differences were assessed with linear regression, and longitudinal analysis of child immune trajectories was evaluated with linear mixed models. Latent cytokine groupings were assessed using an integrated ANOVA framework with principal component analysis.

**Results:** Pregnant women living with HIV had lower GM-CSF, IL-10, IL-12p70, IL-13, IL-2, IL-4, IL-6, IL-7, NGAL, and MMP-9 levels, and higher TNF-α, IFN-γ, and sCD14 levels compared to women without HIV. HEU children had lower GM-CSF, IL-10, IL-12p70, IL-1β, IL-2, and IL-4 and higher sCD14 levels over time compared to HUU children and revealed distinct immunoregulatory profiles. ART initiation during compared to before pregnancy was associated with higher immune marker levels in mothers but not in their children.

**Conclusions:** Altered immune responses are present in mothers with HIV that persist longitudinally in their HEU children, potentially contributing to their health outcomes.

## BACKGROUND

Sub-Saharan Africa has made substantial progress in reducing the risk of vertical HIV [1], leading to a growing number of children who are HIV-exposed uninfected (HEU) [2]. There are over 15 million HEU children worldwide, with approximately 90% in sub-Saharan Africa [3, 4]. South Africa has the highest number of people living with HIV globally, and accounts for approximately 25% of the global population of children who are HEU [5].

Children who are HEU are at increased risk of poorer health, development, and survival outcomes, including immunological vulnerabilities, and lower birth weight compared to HIV-unexposed children (HUU) [1, 6, 7]. The mechanisms through which HIV exposure influences gestation and child outcomes [8], remain incompletely understood. Growing evidence suggests that a dysregulated immune system during pregnancy and in early childhood may contribute to the poorer health of HEU children [8–14]. Furthermore, antiretroviral treatment (ART) exposure during pregnancy may influence the immune regulation of children [15, 16].

Cytokines play an important immunomodulatory function throughout pregnancy in regulating the immune equilibrium to promote foetal growth while allowing maternal immune defences to potential pathogens [17]. The immune system shifts from T helper 1 (Th1) to T helper 2 (Th2) dominance, reducing excessive pro-inflammatory activity and reducing the risk of foetal rejection and miscarriage [18]. HIV infection can disrupt maternal cytokine levels [13, 19–21], and aberrant immune regulation has been reported in HEU children [10, 13]. Studies, including the Drakenstein Child Health Study (DCHS), found cytokine and activation marker differences in HEU children during their first two years [11, 13, 22]. However, longitudinal studies on immune regulation in early childhood among African HEU children are lacking.

This study seeks to understand HIV’s impact on early-life immune regulation in mothers and children, building on our previous findings [13], of lower pro- and anti-inflammatory cytokine levels in mothers and their HEU children. Additionally, a characteristic of HIV infection is monocyte activation, and the findings suggested an increase in monocyte activation during early life. We hypothesised that pregnant mothers living with HIV and their HEU children would have lower cytokine levels than mothers without HIV and their HUU children. The study’s aims were: (1) to cross-sectionally compare cytokines in pregnant women with and without HIV; (2) to determine longitudinal differences in cytokines between HEU and HUU children; (3) to longitudinally evaluate the latent grouping of child cytokines separately in HEU and HUU children. Secondary analysis examined the association of prenatal ART initiation (before or during pregnancy) on cytokine levels.

## METHODOLOGY

### Site

This study was nested in the DCHS, a multidisciplinary, population-based birth cohort. The Drakenstein sub-district, 60 km from Cape Town, has about 300,000 residents. Pregnant women in their 20th to 28th week of gestation, receiving antenatal care, were enrolled between 2012 and 2015 at either TC Newman Clinic, which serves a population of mixed ancestry, or Mbekweni Clinic, which serves a population of black African descent in the Drakenstein sub-district [23, 24]

### Study participants

Pregnant women (n = 1137) recruited for the DCHS had to be at least 18 years old, provide written, signed consent, and have plans to remain in the area for at least 1 year to be eligible for enrolment. Of these, 399 participants (n=179 mothers with HIV and n=220 without HIV) who had available antenatal blood samples were included in this sub-study. Mothers with and without HIV were selected based on demographics and sample availability at different child age points. The DCHS and this sub-study were approved by the University of Cape Town’s HREC (HREC 401/2009 and HREC 053/2023), Stellenbosch University (N12/02/0002), and the Western Cape Provincial Health Research Committee (2011RP45). Mothers provided written informed consent for their enrolment and were reconsented annually in English, Afrikaans, or isiXhosa, depending on the mother’s preference.

### Measurements of maternal demographics and health

Prenatal assessments occurred in the second trimester at the primary healthcare clinics [24]. Sociodemographic data were gathered through an interviewer-administered questionnaire adapted from the South African Stress and Health Study. The mother’s education, household income, and employment status were self-reported. Sociodemographic measures generated a socioeconomic status (SES) score, which was categorised as low, low-to-moderate, moderate-to-high, and high for comparison [23]. Maternal body mass index (BMI) was measured at enrolment. Data on maternal ART use, infant prophylaxis, maternal CD4 count, and viral load were obtained from clinical notes, interviews, and the National Health Laboratory Service system. HIV parameters included maternal CD4+ count during pregnancy and viral load categorised as below the detectable limit (<40 copies/mL), detectable (≥40-1000 copies/mL), and unsuppressed (>1000 copies/mL). ART initiation was categorised as ‘before pregnancy’ or ‘during pregnancy’ [25]. In accordance with the Western Cape government’s guidelines, all mothers living with HIV were initiated on option B+ ART [14, 25]. Prenatal alcohol use was assessed using the Alcohol, Smoking, and Substance Involvement Screening Test (ASSIST) and classified as moderate-severe exposure vs. unexposed [23]. Maternal smoking was categorised based on urine cotinine: non-smokers (<10 ng/ml), passive (10-500 ng/ml), active (>500 ng/ml) [26].

### Postnatal measures

Trained staff collected detailed birth data, including delivery mode, gestational age, sex, head circumference, length, and weight. Gestational age was calculated using the most accurate anticipated delivery date derived from prenatal ultrasound, or if unavailable, the symphysis-fundal height or the last menstrual cycle; prematurity was defined as <37 weeks gestation [27]. HEU children’s HIV-negative status was confirmed at 6 weeks by PCR and at 9 and 18 months by ELISA or after breastfeeding cessation [28]. All HEU infants received prophylaxis as per guidelines (nevirapine only or with zidovudine) from birth [25].

### Immune Assays

Blood serum samples from mothers antenatally and from the children at 6 weeks, 2, 3, and 5 years were analysed for granulocyte-macrophage colony-stimulating factor (GM-CSF), interferon-γ (IFN-γ), interleukin (IL)-1β, IL-2, IL-4, IL-5, IL-6, IL-7, IL-8, IL-10, IL-12p70, IL-13, and tumour necrosis factor-α (TNF-α) concentrations with a Milliplex® premix 13-plex kit (HSTCMAG28SPMX13; Merck) according to the manufacturer’s instructions on a Luminex system (Bio-Plex 200 System; Bio-Rad). Soluble cluster of differentiation (sCD)14, sCD163, chitinase-3-like protein 1 (YKL-40), matrix metalloproteinase-9 (MMP-9), and neutrophil gelatinase-associated lipocalin (NGAL) were measured with enzyme-linked immunosorbent assay (ELISA) kits in accordance with the manufacturer’s instructions (R&D Systems, Minneapolis, USA). All samples were assayed in duplicate and blinded for participant data.

### Statistical analysis

Statistical analyses were conducted using SPSS (version 29, IBM, USA) and R (version 4.2.1, 2022). Data normality was assessed, leading to log-transformation for non-normally distributed markers. Values of cytokine levels below the detection threshold, GM-CSF (n = 9), INF-γ (n = 2), IL-1β (n = 25), IL-2 (n = 4), IL-5 (n = 13), IL-6 (n = 5), IL-4 (n = 5), IL-10 (n = 3), IL-12p70 (n = 3), and IL-13 (n = 17), were substituted with half the minimum detectable value from the standard curve to minimize statistical bias. The Benjamini-Hochberg (BH) was used for multiple comparisons correction. Though the 5% cut-off value is used to refer to statistical significance, observed p-values are reported to indicate the relative strength of associations.

Covariates were selected a priori based on their relevance to inflammatory markers. The unadjusted model included maternal sociodemographic factors (age, alcohol use, BMI, and SES), while model 2 added infant health factors (gestational age and child sex). Continuous variables between HEU and HUU groups were compared using the Wilcoxon test, and Pearson’s chi-squared test evaluated categorical variable associations. Multiple linear regression assessed maternal cytokines by HIV status, adjusting for covariates. Linear mixed models analysed the relationship between maternal HIV status and child cytokine levels over time, using log-transformed cytokine values as dependent variables with random intercepts and slopes. The Akaike information criterion indicated that a variance-components covariance structure best explained the cytokine-predictor relationships, with covariate adjustments.

To evaluate latent child cytokine groupings longitudinally in HEU and HUU children, we used ANOVA simultaneous component analysis (ASCA), extended for repeated measures (RM-ASCA+), which combines linear mixed models with PCA +. It involved three steps: 1. Fitting a mixed-effect linear regression for 18 cytokine responses at 4 time points across n participants, with a design matrix for measurement time, HIV exposure, and their interaction. Time was treated as a categorical variable with 4 levels, and we restricted the random effects to subject-specific intercepts. 2. We decomposed the fixed effect component of the model into effect matrices related to the time and HIV × time components. Principal component analysis (PCA) was then applied to these effect components. 3. Resampling methods were utilised to prevent overfitting and to assess the confidence of the estimated scores and loadings. This method identified underlying immune marker profiles over time for the two groups and estimated the loadings of 18 cytokines on these components, and estimated their associated PCA scores for further analyses.

## RESULTS

### Characteristics of study participants

*Table 1* shows mothers without HIV (n=220) had a lower mean age (25.9 years) compared to those with HIV (n=179) (29.6 years; p<0.001). Mbekweni clinic had a higher prevalence of mothers living with HIV, compared to TC Newman clinic (p<0.001). There were significant differences (p=0.04) in mothers’ education levels by HIV status at all levels; most (n=125) had some secondary education. Mothers without HIV had lower BMI than those with HIV (p=0.01).

**Table 1.** Descriptive demographic characteristics for HIV-infected and HIV-uninfected mothers and neonatal measures at birth.

| **MOTHERS** | **HIV-uninfected** | **HIV-infected** | **P-value** |
| --- | --- | --- | --- |
| **N (%)** | 220 (55.1) | 179 (44.9) | - |
| **Age, mean (SD)** | 25.9 (5.8) | 29.6 (5.2) | <0.001 |
| **Site, N (%)** |  |  | <0.001 |
| *Mbekweni* | 86 (39.1) | 165 (92.2) |  |
| *TC Newman* | 134 (60.9) | 14 (7.8) |  |
| **Education, N (%)** |  |  | 0.04 |
| *Primary* | 14 (6.4) | 18 (10.1) |  |
| *Some secondary* | 125 (56.8) | 115 (64.2) |  |
| *Completed secondary* | 66 (30.0) | 42 (23.5) |  |
| *Any tertiary* | 15 (6.8) | 4 (2.2) |  |
| **Married, N (%)** |  |  | 0.65 |
| *Married/cohabitating* | 88 (40.2) | 76 (42.5) |  |
| *No partner* | 131 (59.8) | 103 (57.5) |  |
| **Body Mass Index (BMI), mean (SD)** | 28.2 (6.67) | 29.9 (6.29) | 0.01 |
| **Smoking (urine cotinine levels ng/ml), N (%)** |  |  | <0.001 |
| *Non-smoker* | 47 (21.4) | 48 (26.8) |  |
| *Passive smoker (10-500 ng/ml)* | 87 (39.5) | 83 (46.4) |  |
| *Active smoker (>500 ng/ml)* | 86 (39.1) | 48 (26.8) |  |
| **Alcohol exposure, N (%)** | 37 (17.5) | 14 (9.3) | 0.03 |
| **Preeclampsia, N (%)** | 5 (2.3) | 3 (1.6) | 0.67 |
| **Maternal CD4+ count in pregnancy, mean (SD)** |  | 478.76 (229.79) | - |
| **Viral load during pregnancy, N (%)** |  |  |  |
| *Below detectable limit (<40 copies/mL)* |  | 71 (39.7) | - |
| *Detectable (≥40-1000 copies/mL)* |  | 25 (14.0) | - |
| *Unsuppressed (>1000 copies/mL)* |  | 14 (7.8) | - |
| **Antiretroviral regimen during pregnancy, N** |  |  |  |
| **(%)** |  |  |  |
| *Prevention of mother-to-child transmission prophylaxis/ AZT monotherapy* | - | 26 (14.5) | - |
| *First-line triple therapy (NNRTI + 2NRTIs)* | - | 141 (78.8) | - |
| *Second-line (PI-containing)* | - | 10 (5.6) | - |
| **Initiation of antiretroviral treatment, N (%)** |  |  |  |
| *Before pregnancy* | - | 74 (41.3) | - |
| *During pregnancy* | - | 103 (57.5) | - |
| **Social-economic status (SES), N (%)** |  |  | 0.18 |
| *Lowest SES* | 54 (24.5) | 60 (33.5) |  |
| *Low-moderate SES* | 52 (23.6) | 41 (22.9) |  |
| *Moderate-high SES* | 63 (28.6) | 48 (26.8) |  |
| *High SES* | 51 (23.2) | 30 (16.8) |  |
| **Delivery Method, N (%)** |  |  | 0.04 |
| *Vaginal* | 183 (83.6) | 134 (75.3) |  |
| *Elective Caesarean* | 36 (16.4) | 44 (24.7) |  |
| **INFANTS AT 6-10 weeks** | **HUU** | **HEU** | **P-value** |
| **N (%)** | 169 (56.3) | 131 (43.7) | - |
| **Age (months), mean (SD)** | 1.79(0.32) | 1.88 (0.43) | 0.05 |
| **Sex (Female) N (%)** | 71 (55.5) | 57 (44.5) | 0.80 |
| **Prematurity (<37 weeks), N (%)** | 25 (58.1) | 18 (41.9) | 0.80 |
| **Birth weight (Kg), mean (SD)** | 3.032 (554.28) | 2.999 (558.74) | 0.61 |
| **CHILDREN 2 YEARS** | **HUU** | **HEU** | **P-value** |
| **N (%)** | 195 (62.9) | 115 (37.1) | - |
| **Age (months), mean (SD)** | 25.1 (0.79) | 25.1 (0.88) | 0.70 |
| **Sex (Female) N (%)** | 82 (61.2) | 52 (38.8) | 0.68 |
| **Prematurity (<37 weeks), N (%)** | 26 (66.7) | 13 (33.3) | 0.70 |
| **Birth weight (Kg), mean (SD)** | 3.024 (586.35) | 3.039 (525.46) | 0.83 |

| CHILDREN 3 YEARS | HUU | HEU | P-value |
| --- | --- | --- | --- |
| N (%) | 153 (60.5) | 100 (39.5) | - |
| Age (months), mean (SD) | 37.4 (0.86) | 37.2 (0.66) | 0.21 |
| Sex (Female) N (%) | 61 (58.7) | 43 (41.3) | 0.62 |
| Prematurity (<37 weeks), N (%) | 18 (58.1) | 13 (41.9) | 0.77 |
| Birth weight (Kg), mean (SD) | 3.035 (579.29) | 3.049 (577.71) | 0.85 |

| CHILDREN 5 YEARS | HUU | HEU | P-value |
| --- | --- | --- | --- |
| N (%) | 160 (59.3) | 110 (40.7) | - |
| Age (months), mean (SD) | 61.6 (0.83) | 61.8 (0.92) | 0.33 |
| Sex (Female) N (%) | 67 (57.8) | 49 (42.2) | 0.66 |
| Prematurity (<37 weeks), N (%) | 20 (60.6) | 13 (39.4) | 0.87 |
| Birth weight (Kg), mean (SD) | 3.049 (594.22) | 3.009 (557.72) | 0.58 |

**Abbreviations:** SD, standard deviation; N, numbers; AZT, zidovudine; NNRTI, nucleoside reverse transcriptase inhibitors; 2NRTIs, two nucleoside reverse transcriptase inhibitors; PI, protease inhibitor

More active smokers were among mothers without HIV (p<0.001). Vaginal births were more common in mothers without HIV (n=183), while caesarean sections were higher in mothers with HIV (n=44) (p=0.04). Fewer mothers living with HIV were exposed to alcohol compared to mothers without HIV (p=0.03). Immune marker data at each time-point and the number of repeated measures are depicted in *Supplementary Table 1*.

### Cytokine levels in pregnant women with and without HIV

*Figures 1A-R* illustrate differences in cytokines between pregnant women with and without HIV after adjusting for covariates. GM-CSF (*figure 1A*), IL-10 (*figure 1C*) IL-12p70 (*figure 1D*), IL-13 (*figure 1E*), IL-2 (*figure 1G*), IL-4 (*figure 1H*), IL-6 (*figure 1J*), IL-7 (*figure 1K*), NGAL (*figure 1N*), and MMP-9 (*figure 1O*) levels were significantly lower in women with HIV, while IFN-γ (*figure 1B*), TNF-α (*figure 1M*) and sCD14 (*figure 1R*) were higher (all p ≤ 0.05). However, after the BH correction for multiple comparisons, GM-CSF, IL-10, IL-12p70, IL-13, IL-2, IL-4, IL-6, NGAL, MMP-9, and IFN-γ remained significant. Additional regression analysis outcomes are provided in *Supplementary Data Table 2*.

**Figure 1.**
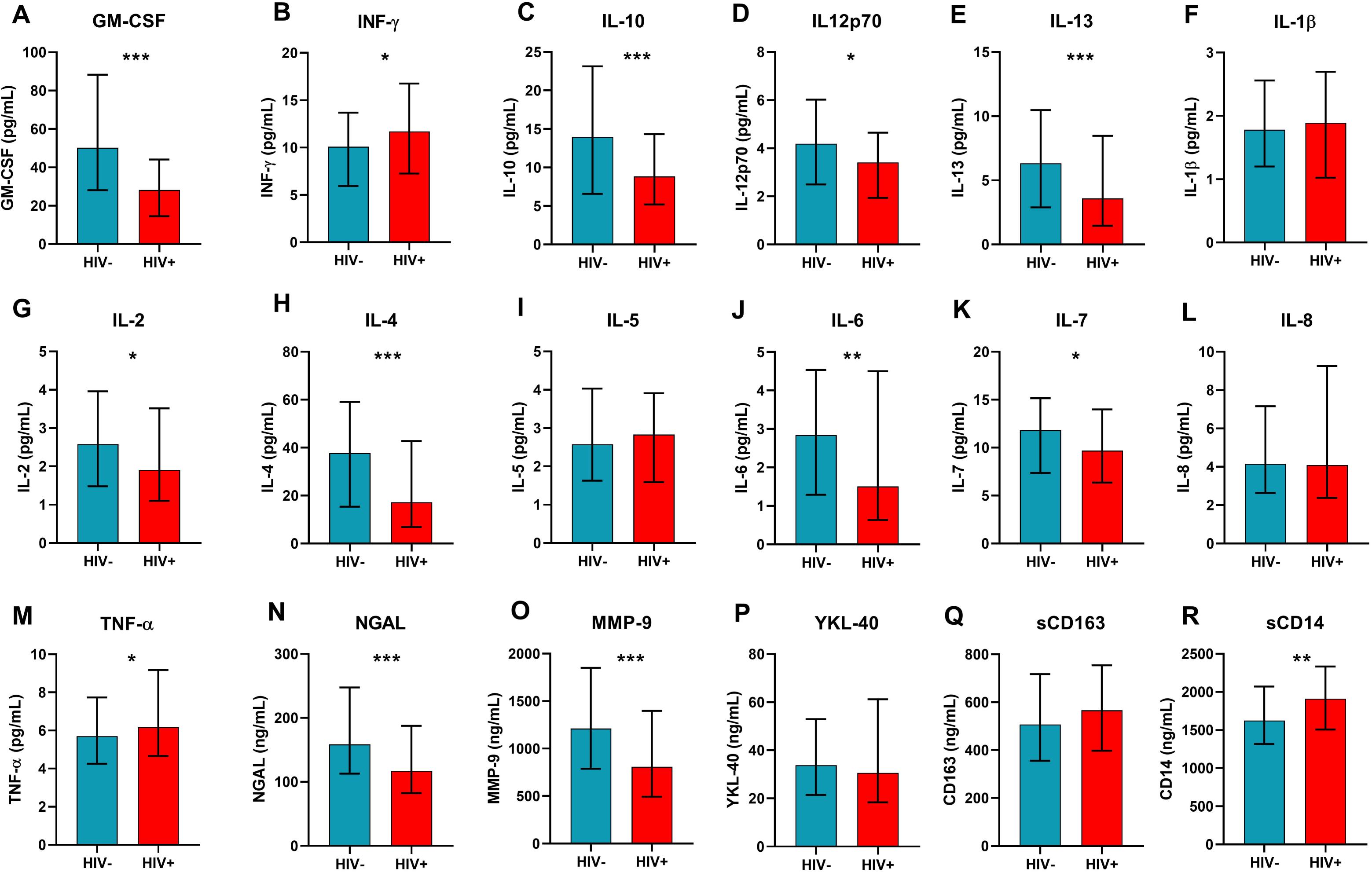
Cytokines during pregnancy by maternal HIV status. Significant values are adjusted for covariates. Bars represent median protein concentrations in the study groups, with error bars indicating the interquartile ranges. *p<0.05, **p<0.01, ***p<0.001. Abbreviations: Granulocyte-macrophage colony-stimulating factor, GM-CSF; interferon-γ, IFN-γ; <u>interleukin</u>, IL; tumour necrosis factor-α, TNF-α; neutrophil gelatinase-associated <u>lipocalin</u>, NGAL; metalloproteinase-9, MMP-9; soluble cluster of differentiation, sCD; chitinase-3-like protein 1, YKL-40. **Alt text:** Bar graphs illustrating median cytokine concentrations during pregnancy among mothers with and without HIV, with error bars representing interquartile ranges and statistically significant differences marked by asterisks.

### Longitudinal measures of inflammatory markers in HUU and HEU children

*Figures 2A-R* illustrate longitudinal cytokine levels over the first 5 years in HEU compared to HUU children. In the unadjusted linear mixed models (*Figure 2S*) GM-CSF (coefficient = 0.221, 95% CI =-0.358 to -0.085, p = 0.002), IL-10 (coefficient = -0.215, 95% CI = 0.312 to -0.118, p = 0.000), IL-12p70 (coefficient = -0.118, 95% CI = [-0.216 to -0.021, p = 0.010), IL-2 (coefficient = -0.127, 95% CI = -0.235 to -0.018, p = 0.006) and IL-4 (coefficient = -0.193, 95% CI = -0.364 to -0.021, p = 0.003) were significantly lower over time in HEU compared to HUU children, and sCD14 (coefficient = 0.078, 95% CI = 0.031 to 0.124, p = 0.001) was significantly higher.

**Figure 2.**
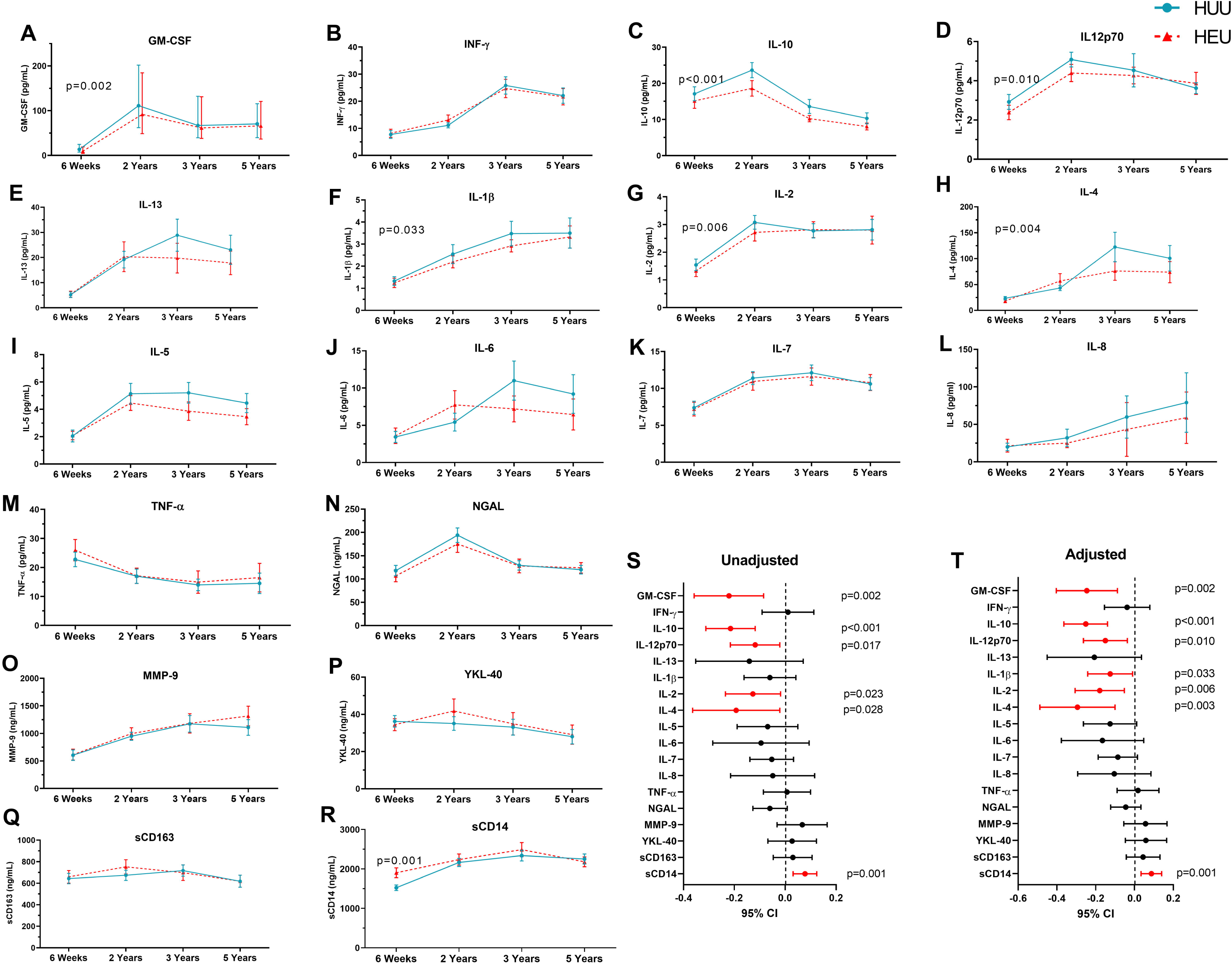
Longitudinal differences in cytokines between HEU and HUU children. Bars indicate median protein concentrations in the study groups, and error bars represent the interquartile ranges. Abbreviations: Granulocyte-macrophage colony-stimulating factor, GM-CSF; interferon-γ, IFN-γ; <u>interleukin</u>, IL; tumour necrosis factor-α, TNF-α; neutrophil gelatinase-associated <u>lipocalin</u>, NGAL; metalloproteinase-9, MMP-9; soluble cluster of differentiation, sCD; chitinase-3-like protein 1, YKL-40. **Alt text:** Line graphs (labelled A to F) and forest plots (labelled S and T). comparing longitudinal differences between HEU and HUU children, with bars representing protein median concentrations and error bars representing interquartile ranges.

After adjusting for covariates (*Figure 2T*) GM-CSF (coefficient = -0.246, 95% CI =-0.403 to - 0.088, p = 0.002), IL-10 (coefficient = -0.251, 95% CI = -0.364 to -0.139, p < 0.001), IL-12p70 (coefficient = -0.150, 95% CI = -0.263 to -0.037, p = 0.010), IL-1β (coefficient = -0.125, 95% CI = -0.241 to -0.010, p = 0.033), IL-2 (coefficient = -0.179, 95% CI = -0.306 to -0.052, p = 0.006) and IL-4 (coefficient = -0.294, 95% CI = -0.488 to -0.100, p = 0.003) remained significantly lower over time and sCD14 (coefficient = 0.087, 95% CI = 0.034 to 0.140, p = 0.001) higher in HEU children as compared to HUU. After BH correction, GM-CSF, IL-10, IL-12p70, IL-2, and IL-4 were significantly lower, while sCD14 was higher in HEU children. A summary is provided in *Supplementary Table 3*. Correlograms visualised cytokine relationships between mothers and children according to maternal HIV status (*supplementary data figure 1*). Correlations observed in the first two years diminished by 3 and 5 years. *Supplementary Table 4* further summarises regression coefficients and cytokine levels in children.

### Principal Component Analysis of Time and HIV Effects on Cytokine Profiles

*Figure 3* illustrates the PCA of the effects of time on cytokine levels in HUU and HEU children. *Figure 3A* reveals that PCA1 accounted for 81% of the variability with increasing levels from 6 weeks to 2 years and a stable profile from 2 to 5 years in children who are HUU and HEU. GM-CSF, IFN-γ, IL-1β, IL-2, IL-5, IL-3, IL-12p70, IL-4, MMP-9, sCD14, and IL-7 were positively associated with this component, while TNF-α was negatively associated with it. *Figure 3B:* PCA2 explained 15% of the variability, showing a profile that increased from 6 weeks to 2 years and a decreased profile from 2 to 5 years. IL-10 and NGAL exhibit a strong positive association with this profile, while IFN-γ and IL-1β demonstrate a strong negative association. Both PCA1 and PCA2 profiles indicate lower scores for the HEU group at earlier times, with the difference between the two HIV-exposure groups becoming smaller over time.

**Figure 3.**
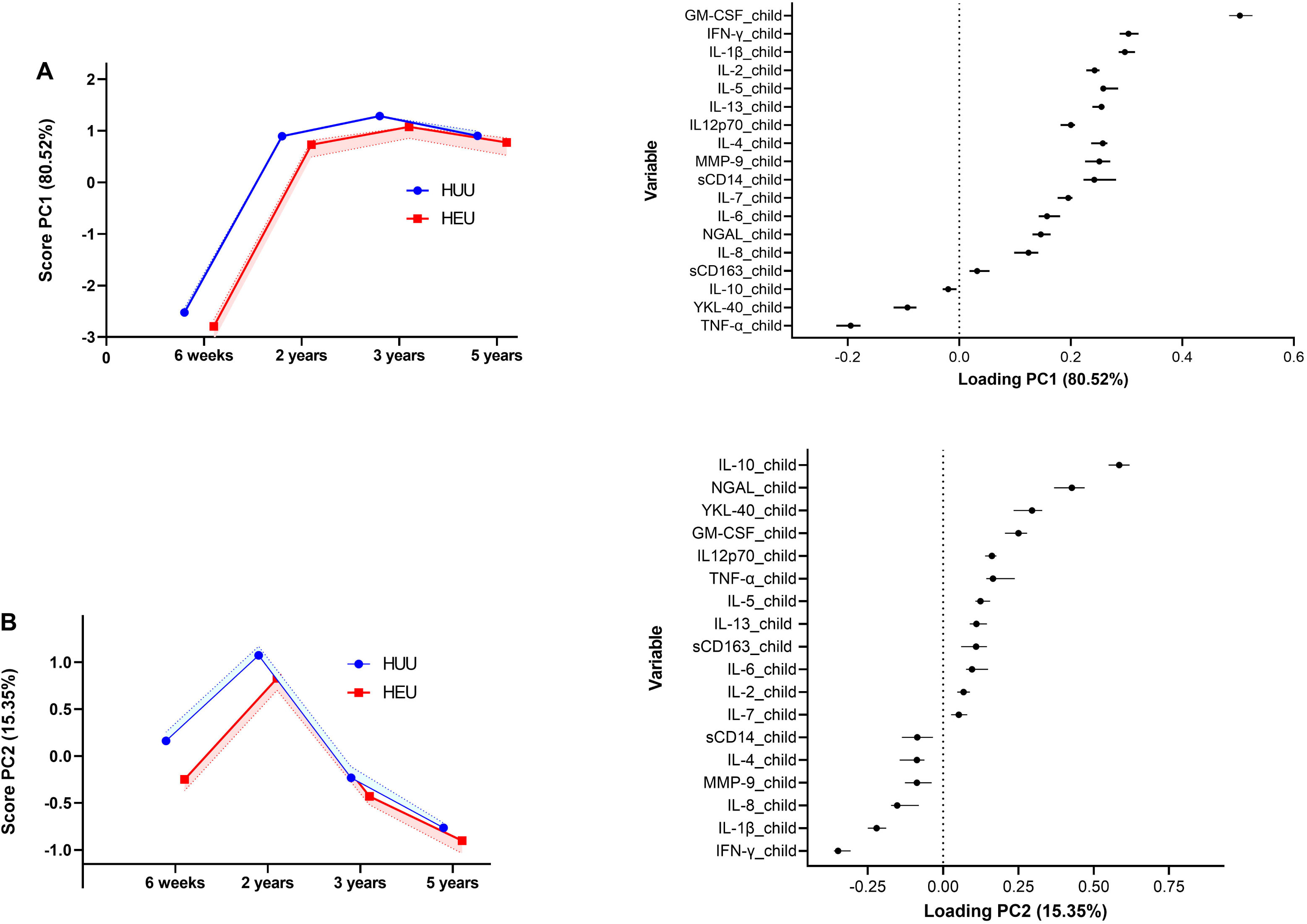
An illustration of the principal component analysis (PCA), with **(A)** PCA1 accounting for 80.5% of the variability and reflecting an increasing profile from 6 weeks to 2 years, and **(B)** PCA2 accounting for 15.35% of the variability and reflecting a profile that first increased to 2 years, whereafter it decreased at each time point. **Alt text:** Graphs on the left show PCA scores over time for HUU (blue line) and HEU (green line) children, while the graphs on the right illustrate trends of PCA loadings.

*Figure 4* shows cytokine profile differences over time in HEU children compared to HUU children. PCA1 *(Figure 4A)* highlights differences at 6 weeks that diminish by 5 years. sCD14 exhibits a strong positive association with this component, while IL-12p70, NGAL, IL-10, IL-2, IL-4, and IL-1β are negatively associated with it. In *Figure 4B,* PCA2 shows no early differences; however, the profiles diverge at later time points. Cytokines IL-5, IL-10, IL-6, IL-4, and IL-13 were positively associated with PCA2, while MMP-9 was negatively associated with it.

**Figure 4.**
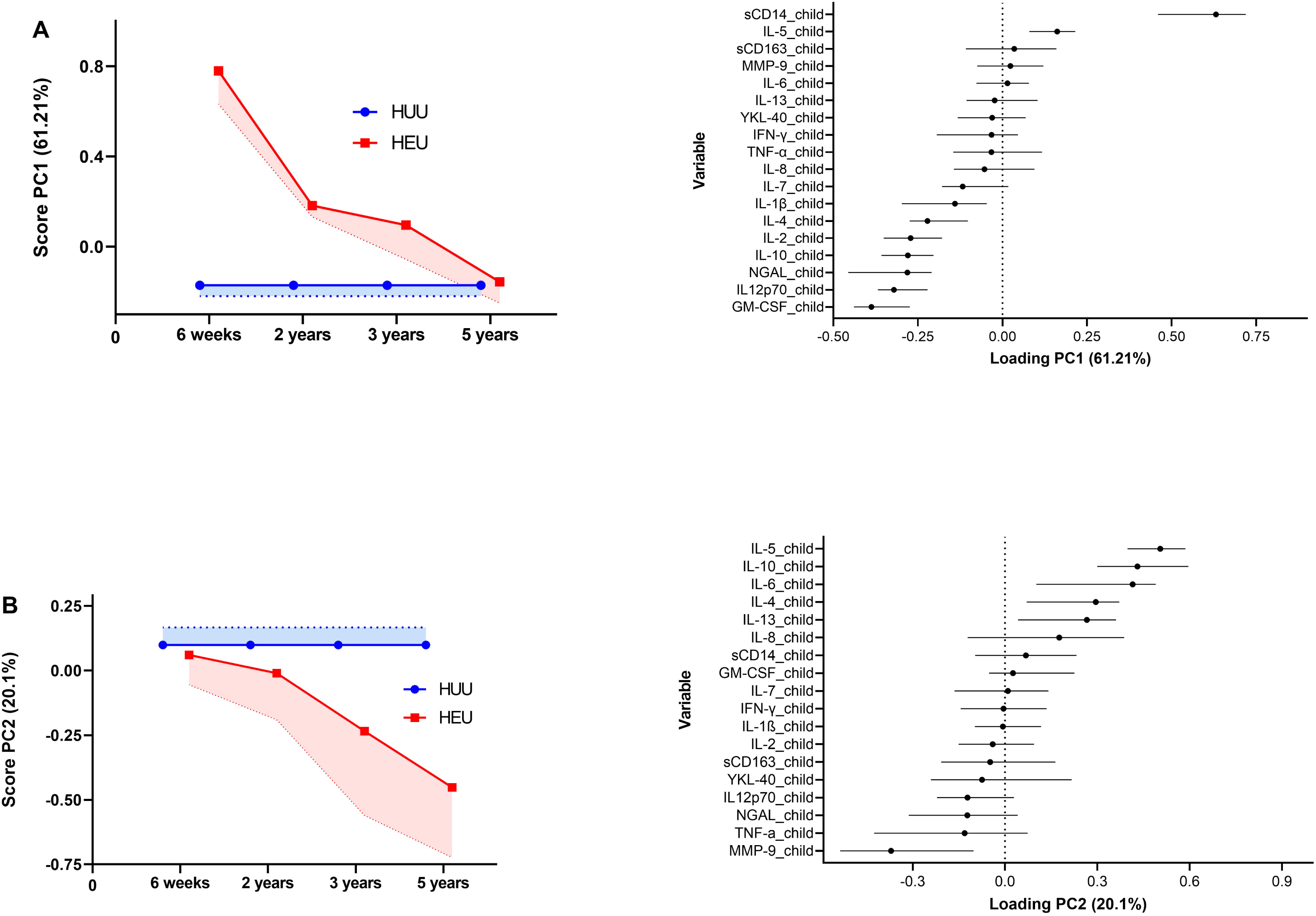
An illustration of the first two Principal component analysis (PCA) components relevant to the HIV-exposure groups, with the HIV-unexposed serving as the reference category; **(A)** PCA1 represents a component that captures a significant difference between the HIV-exposed and unexposed groups at earlier time points, which subsequently diminishes and converges at a later time point. **(B)** PCA2 shows no difference between the two HIV-exposure groups early on, but their profiles diverge at later time points. **Alt text:** Graphs on the left show PCA scores over time for HUU (blue line) and HEU (green line) children, while the graphs on the right illustrate trends of PCA loadings.

### Evaluating the relationship between maternal ART initiation and cytokine levels

Maternal ART initiation was associated with higher cytokine levels of GM-CSF, IFN-γ, IL- 12p70, IL-13, IL-2, IL-4, and IL-5 (all p>0.05) in mothers who initiated treatment during pregnancy versus those who received treatment before pregnancy (*Figure 5*). After BH adjustment, GM-CSF and CD14 remained significant. However, no significant differences in cytokine levels were observed in HEU children whose mothers’ received treatment during pregnancy compared to before pregnancy (*Supplementary Table 5*).

**Figure 5.**
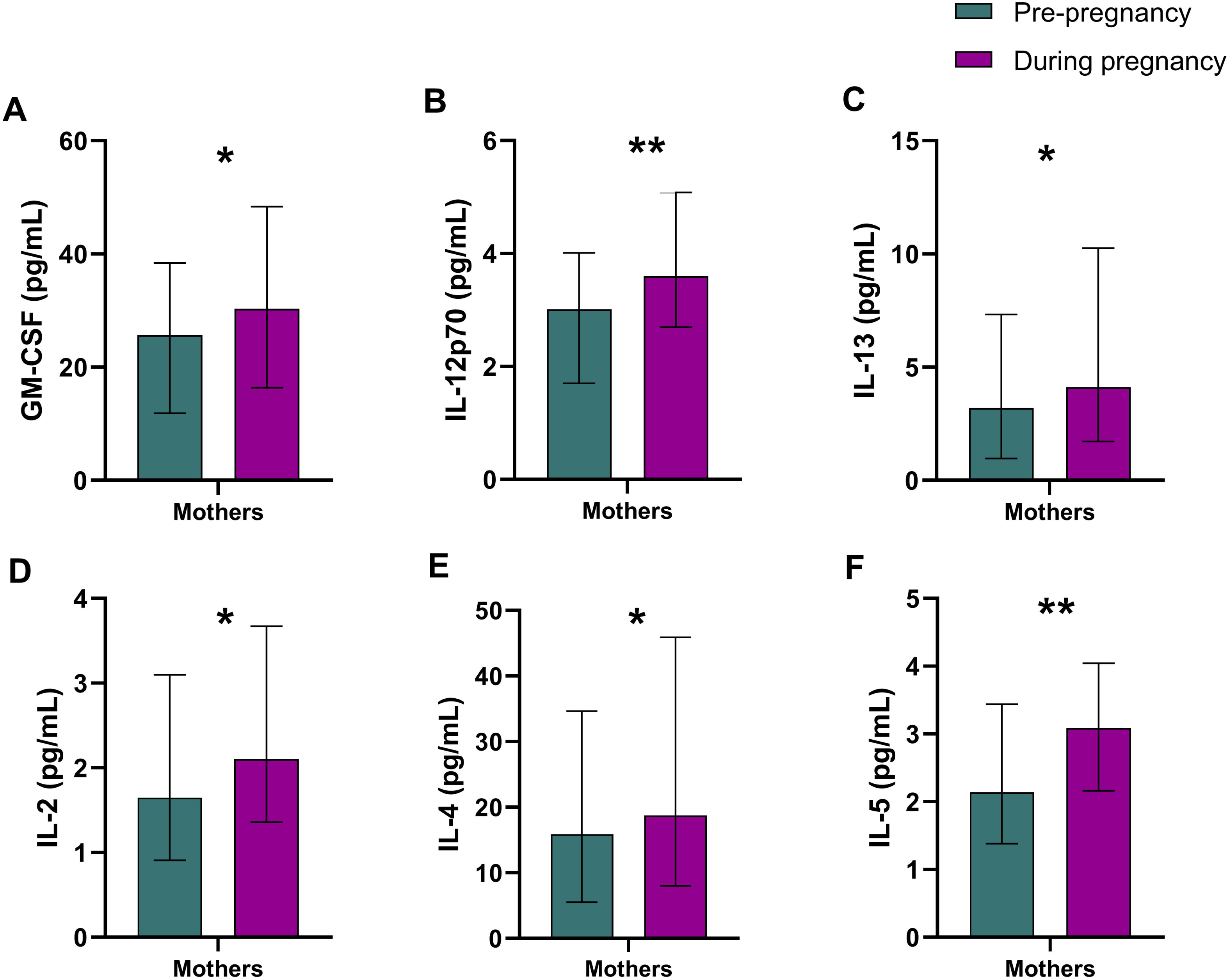
Illustration of the immunological effect of ART initiation pre-pregnancy and during pregnancy. Significant values are adjusted for covariates. Error bars indicating the interquartile ranges. *p<0.05, **p<0.01. Abbreviations: Granulocyte-macrophage colony-stimulating factor, GM-CSF; <u>interleukin</u>, IL. **Alt text:** Bar graphs comparing ART initiation before and during pregnancy with statistically significant differences being marked by asterisks and error bars representing interquartile ranges.

## DISCUSSION

This study found that South African women living with HIV have dysregulated cytokine levels during pregnancy with lower GM-CSF, IL-12p70, IL-13, IL-2, IL-4, IL-6, IL-7, IL-10, NGAL, and MMP-9 levels and higher IFN-γ, TNF-α, and sCD14 levels than women without HIV. Similarly, HEU children showed lower GM-CSF, IL-10, IL-12p70, IL-1β, and IL-2, with higher sCD14 levels in the first five years of life. Longitudinal PCA revealed distinct patterns in immunoregulatory profiles of HEU children during the first two years of life and from 2 years to 5 years of age. Initiating ART during pregnancy was associated with higher cytokine levels in mothers living with HIV compared to initiation before pregnancy.

This study highlights immune dysregulation in pregnant women living with HIV. Overall, our results demonstrate significantly reduced levels of both pro- and anti-inflammatory cytokines among pregnant women living with HIV compared to those without, aligning with previous research [13, 21, 29–31]. Some studies reported high inflammatory cytokines in pregnant women with HIV [19, 32]. Our findings indicate a disrupted Th1/Th2 balance, with heightened Th1 and diminished Th2 cytokine environment compared to HIV-negative pregnant women. Elevated Th1 cytokines during pregnancy have been associated with complications like pre- eclampsia and preterm delivery [33]. Consistent with our findings, elevated levels of IFN-γ [34], TNF-α [35, 36] and sCD14 [35] have been reported, whereas lower levels of IFN-γ [37], TNF-α [21] and sCD14 [38] were found in mothers living with HIV across other settings. This may be partly explained by the different subtypes of HIV that are prevalent based on geographical regions, which may be immunosuppressive in the case of subtype C (more prevalent in South Africa) [39].

In children who are HEU, this study corroborates our previous findings, which reported reduced levels of inflammatory cytokines, including IFN-γ and IL-1β, at 6 weeks, as well as lower levels of IFN-γ, IL-1β, IL-2, and IL-4 at 2 years [13]. Consistent with our findings, a recent study documented lower levels of IL-12p70 in Kenyan HEU children aged 18–36 months [40]. Borges-Almeida et al. (2011) also observed reduced IL-4 levels in HEU newborns and elevated IFN-γ levels. In Kenya, a longitudinal study reported non-significantly lower IL-10 and IL-12p70 levels in HEU children compared to HUU children during the first year [22]. Studies show South African children have lower immune responses than those in North America, South America, and Europe [41, 42]. This observation is further supported by our findings, which demonstrate lower cytokine levels in South African HEU children compared to HUU children, as well as other sub-Saharan African populations. In contrast, a study in the United States reported higher cytokine levels (e.g., TNF-α, IL-6, IL-10 in newborns; IFN-γ, TNF-α, IL- 2, IL-1β in 6-month-olds) in HEU infants compared to HUU [36]. Additionally, a Brazilian study found elevated IL-6 levels in HEU children at birth, which remained higher until 6 months [43]. Elevated levels of sCD14 have also been consistently reported in HEU children across various studies [11, 12, 44]. The disparities in cytokine levels among HEU children may partly be attributed to population-specific immune development.

Longitudinally, PCA revealed similar immune profiles over time in both HEU and HUU children. However, HEU children had lower levels of these immune profiles, particularly during the first two years, suggesting that immune maturation is predominantly affected during this critical developmental window. PCA also highlighted elevated sCD14 levels in HEU children, which accounted for the largest differences observed at earlier time points. As HEU children aged, their immune profiles became increasingly distinct, characterised by declining levels of IL-5, IL-10, IL-6, IL-4, and IL-13. Collectively, these findings indicate that HEU children develop distinct immune profiles over the first five years, which may have implications for their long-term health and susceptibility to disease. Further studies are warranted to explore the underlying mechanisms driving these immune alterations and their potential clinical consequences.

The study found altered cytokine levels in pregnant women living with HIV and their HEU children, despite all mothers receiving ART. However, we found that the absence of ART prior to pregnancy was associated with lower cytokine levels compared to pregnant women who had initiated ART before conception. This finding aligns with a study from Mozambique [45], suggesting that pre-pregnancy ART may stabilise cytokine levels. The effects of ART on cytokines during pregnancy remain a subject of debate. Some studies indicate that ART alters cytokine levels [19, 21, 46, 47], while others reported no significant impact [48, 49]. Furthermore, our results showed no significant differences in cytokine levels between ART initiation and the children’s cytokine levels. The impact of the ART environment on maternal health and children’s immune profiles needs further investigation.

We observed weak correlations between cytokine levels in mothers and their children, consistent with a previous finding in HEU children [29]. These correlations diminished over time and became negligible by age 5. HEU children showed altered cytokine profiles, with lower levels and higher sCD14, similar to pregnant women living with HIV. This suggests that *in utero* HIV exposure influences foetal immune programming, with lasting effects on the child’s immune system. Higher IFN-γ and TNF-α levels in mothers living with HIV were not present in their HEU children. This may be explained by the activation of the cGAS-STING (cyclic GMP-AMP synthase - stimulator of interferon genes) pathway in mothers, triggered by cytoplasmic HIV DNA and leading to upregulation of IFN-γ and TNF-α [50]. Mothers and their HEU children had elevated sCD14 levels, a marker indicative of monocyte activation and linked to intestinal dysbiosis and microbial translocation. We speculate that elevated sCD14 levels may result from maternal gut microbiota dysbiosis transferred to the child. This could explain why differences between HEU and HUU children were most notable at 6 weeks and diminished by 5 years as the microbiome matured and influenced by environmental factors.

The strengths of this study were that we (i) evaluated cytokine profiles associated with immunological dysregulation in a South African longitudinal birth cohort, addressing the gap in research on maternal HIV exposure, (ii) assessed multiple early life timepoints. Our limitations included (1) measuring maternal cytokines at only one late second-trimester point, which may not accurately represent the dynamic immune changes that occur during pregnancy. (2) Missing data includes (i) unavailability of child samples at specific time points and (ii) lack of HIV viral load data due to changing guidelines. (3) The study’s findings may be influenced by regional or ethnic differences, such as genetics and socioeconomic status, making them primarily relevant to the South African population and not fully generalisable to other global populations. Future recommended work would be to evaluate the association of cytokine profiles with clinical outcomes.

In conclusion, this study reports altered cytokine levels in mothers living with HIV, with a similar immune profile in their HEU children, that persists up to 5 years of age. These findings urge further research into the association between clinical outcomes and immune dysregulation to understand if these may underlie clinical vulnerabilities observed in children who are HEU.

## Author Contributions

**TSK:** Writing – original draft, review & editing, Methodology, Investigation, Visualisation. **CJW:** Visualisation, Writing – review & editing, **JFA:** Methodology, Investigation, Writing – review & editing. **FL:** Formal analysis, Software, Data Curation, Software. **HJZ:** Writing – review & editing, Conceptualisation, Resources. **DJS:** Writing – review & editing, Supervision, Conceptualisation, Resources. **PJWN:** Writing – original draft, review & editing, Conceptualisation, Supervision, Resources, Methodology, Visualisation, Data Curation, Funding acquisition.

## Acknowledgments

We would like to thank and recognise the DCHS staff: the teams responsible for the study data, laboratory, and administrative work. We would like to thank the clinical and administrative staff of the Western Cape Government Department of Health at Paarl Hospital, and the two clinics that were mainly used in this study. We would also like to thank and acknowledge the families and children who took part in this study.

## Data Availability

Due to ethical constraints that every person involved in the study is obligated to meet, data cannot be made publicly available per the DCHS cohort guidelines. The de-identified data used to generate the study’s conclusions can be accessed upon reasonable request to the private investigator involved in this study through the corresponding author.

