## Supplementary Data for "Maternal and early-life longitudinal cytokine profiles in a South African birth cohort: the impact of HIV"

**SUPPLEMENTARY INFORMATION:**

**Supplementary Table 1.** Numbers of available immune data at each time point and numbers of repeated measures

| | Mother | 7 weeks | 2 Year | 3 Year | 5 Year |
|---|---|---|---|---|---|
| **Number (%)** | 399 (100) | 300 (75.2) | 310 (77.7) | 253 (63.4) | 270 (67.7) |
| | One measure | Two measures | Three measures | Four measures | |
| **Numbers of repeated measures (%)** | 53 (13.3) | 92 (23.1) | 120 (30.1) | 134 (33.6) | |

**Supplementary Table 2.** Regression coefficients according to maternal HIV

| Mothers | Unadjusted | | | | Adjusted | | | |
|---|---|---|---|---|---|---|---|---|
| | B (SE) | β | p | BH | B (SE) | β | p | BH |
| **GM-CSF** | -0.614 (0.092) | -0.311 | 0.000 | 0.000 | -0.531(0.098) | -0.27 | 0.000 | 0.000 |
| **IFN-γ** | 0.135 (0.072) | 0.092 | 0.062 | 0.086 | 0.038 (0.077) | 0.113 | 0.032 | 0.048 |
| **IL-10** | -0.326 (0.094) | -0.169 | 0.001 | 0.001 | -0.267(0.100) | -0.139 | 0.008 | 0.017 |
| **IL-12p70** | -0.222 (0.073) | -0.148 | 0.003 | 0.005 | -0.199 (0.078) | -0.133 | 0.011 | 0.021 |
| **IL-13** | -0.523 (0.125) | -0.201 | 0.000 | 0.000 | -0.561 (0.134) | -0.217 | 0.000 | 0.000 |
| **IL-1β** | -0.006 (0.071) | 0.004 | 0.929 | 0.929 | 0.014 (0.076) | -0.241 | 0.033 | 0.850 |
| **IL-2** | -0.268 (0.091) | -0.143 | 0.004 | 0.006 | -0.233 (0.097) | -0.124 | 0.017 | 0.030 |
| **IL-4** | -0.632 (0.127) | -0.237 | 0.000 | 0.000 | -0.625 (0.136) | -0.235 | 0.000 | 0.000 |
| **IL-5** | -0.008 (0.072) | -0.005 | 0.915 | 0.929 | -0.022 (0.077) | -0.013 | 0.779 | 0.850 |
| **IL-6** | -0.400 (0.127) | -0.154 | 0.002 | 0.004 | -0.378 (0.135) | -0.145 | 0.005 | 0.013 |
| **IL-7** | -0.116 (0.056) | -0.101 | 0.039 | 0.058 | -0.123 (0.060) | -0.107 | 0.040 | 0.055 |
| **IL-8** | -0.106 (0.113) | 0.046 | 0.349 | 0.419 | 0.024 (0.121) | 0.011 | 0.840 | 0.850 |
| **TNF-α** | 0.173 (0.055) | 0.148 | 0.002 | 0.005 | 0.135 (0.061) | 0.116 | 0.027 | 0.044 |
| **NGAL** | -0.257 (0.055) | -0.223 | 0.000 | 0.000 | -0.275 (0.059) | -0.244 | 0.000 | 0.000 |
| **MMP-9** | -0.361 (0.068) | -0.252 | 0.000 | 0.000 | -0.387 (0.073) | -0.27 | 0.000 | 0.000 |
| **YKL40** | 0.025 (0.072) | 0.017 | 0.731 | 0.822 | -0.019 (0.077) | -0.014 | 0.802 | 0.850 |
| **sCD163** | 0.073 (0.046) | 0.078 | 0.112 | 0.144 | 0.043 (0.049) | 0.044 | 0.376 | 0.484 |
| **sCD14** | 0.123 (0.034) | 0.173 | 0.000 | 0.001 | 0.109 (0.037) | 0.155 | 0.003 | 0.009 |

Adjusted for age, sex, preterm, socioeconomic status, alcohol use, maternal age, maternal body mass index (BMI), maternal HIV. Abbreviations: Benjamini-Hochberg corrected p-value, BH; Granulocyte-macrophage colony-stimulating factor, GM-CSF; interferon-γ, IFN-γ; interleukin, IL; tumor necrosis factor-α, TNF-α; neutrophil gelatinase-associated lipocalin, NGAL; metalloproteinase-9, MMP-9; soluble cluster of differentiation, sCD; chitinase-3-like protein 1, YKL-40.

**Supplementary Table 3.** Longitudinal cytokine expression in HUU and HEU children

| | Unadjusted | | | | | Adjusted | | | | |
|---|---|---|---|---|---|---|---|---|---|---|
| | Effect | CI | | p | BH | Effect | CI | | p | BH |
| **Markers** | | lower | upper | | | | lower | upper | | |
| **GM-CSF** | -0.221 | -0.358 | -0.085 | 0.002 | 0.012 | -0.246 | -0.403 | -0.088 | 0.002 | 0.012 |
| **IFN-γ** | 0.011 | -0.091 | 0.113 | 0.826 | 0.875 | -0.038 | -0.155 | 0.079 | 0.522 | 0.553 |
| **IL-10** | -0.215 | -0.312 | -0.118 | 0.000 | 0.000 | -0.251 | -0.364 | -0.139 | 0.000 | 0.000 |
| **IL-12p70** | -0.118 | -0.216 | -0.021 | 0.017 | 0.077 | -0.150 | -0.263 | -0.037 | 0.010 | 0.030 |
| **IL-13** | -0.141 | -0.352 | 0.071 | 0.192 | 0.384 | -0.207 | -0.450 | 0.036 | 0.094 | 0.190 |
| **IL-1β** | -0.060 | -0.162 | 0.042 | 0.251 | 0.384 | -0.125 | -0.241 | -0.010 | 0.033 | 0.085 |
| **IL-2** | -0.127 | -0.235 | -0.018 | 0.023 | 0.083 | -0.179 | -0.306 | -0.052 | 0.006 | 0.022 |
| **IL-4** | -0.193 | -0.364 | -0.021 | 0.028 | 0.084 | -0.294 | -0.488 | -0.100 | 0.003 | 0.014 |
| **IL-5** | -0.069 | -0.189 | 0.050 | 0.256 | 0.384 | -0.126 | -0.263 | 0.011 | 0.126 | 0.211 |
| **IL-6** | -0.095 | -0.285 | 0.094 | 0.324 | 0.449 | -0.165 | -0.377 | 0.048 | 0.129 | 0.211 |
| **IL-7** | -0.053 | -0.139 | 0.033 | 0.226 | 0.384 | -0.086 | -0.187 | 0.015 | 0.095 | 0.190 |
| **IL-8** | -0.049 | -0.215 | 0.116 | 0.559 | 0.645 | -0.104 | -0.293 | 0.085 | 0.280 | 0.358 |
| **TNF-α** | 0.007 | -0.086 | 0.100 | 0.876 | 0.876 | 0.018 | -0.090 | 0.126 | 0.747 | 0.747 |
| **NGAL** | -0.060 | -0.127 | 0.008 | 0.083 | 0.213 | -0.045 | -0.123 | 0.033 | 0.257 | 0.358 |
| **MMP-9** | 0.067 | -0.032 | 0.165 | 0.184 | 0.384 | 0.057 | -0.055 | 0.168 | 0.318 | 0.358 |
| **YKL40** | 0.027 | -0.068 | 0.123 | 0.573 | 0.645 | 0.059 | -0.047 | 0.166 | 0.275 | 0.358 |
| **sCD163** | 0.030 | -0.047 | 0.106 | 0.446 | 0.573 | 0.044 | -0.042 | 0.131 | 0.312 | 0.358 |
| **sCD14** | 0.078 | 0.031 | 0.124 | 0.001 | 0.012 | 0.087 | 0.034 | 0.140 | 0.001 | 0.009 |

Adjusted for age, sex, preterm, socioeconomic status, alcohol use, maternal age, maternal body mass index (BMI), maternal HIV. Abbreviations: Benjamini-Hochberg corrected p-value, BH; Granulocyte-macrophage colony-stimulating factor, GM-CSF; interferon-γ, IFN-γ; interleukin, IL; tumor necrosis factor-α, TNF-α; neutrophil gelatinase-associated lipocalin, NGAL; metalloproteinase-9, MMP-9; soluble cluster of differentiation, sCD; chitinase-3-like protein 1, YKL-40.

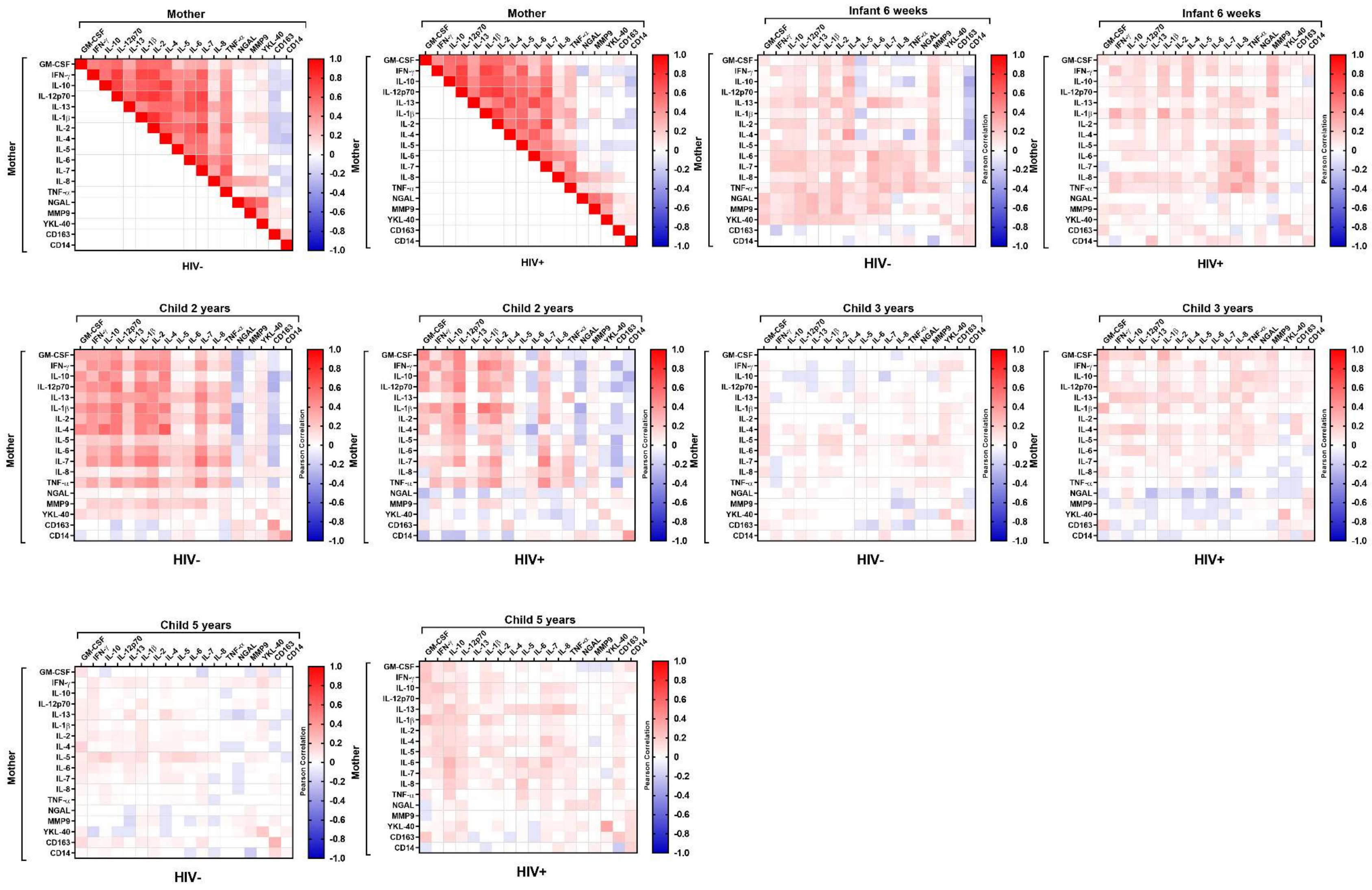

**Supplementary Figure 1. Correlation for mother and child cytokines, stratified according to maternal HIV status.** Granulocyte-macrophage colony-stimulating factor (GM-CSF); interferon-γ (IFN-γ); interleukin (IL); tumor necrosis factor-α (TNF-α); neutrophil gelatinase-associated lipocalin (NGAL); metalloproteinase-9 (MMP-9); soluble cluster of differentiation 14 (sCD14); soluble cluster of differentiation (sCD163); chitinase-3-like protein 1 (YKL-40). **Alt text:** A heatmap displaying correlations for the cytokines of mothers (those living with and without HIV) and their children. Red indicates a positive correlation, while blue indicates a negative correlation.

**Supplementary Table 4**. Regression coefficients for differences in cytokine levels between HEU vs HUU children at each time point.

| | 6 weeks | | | | | | | | 2 years | | | | | | | |
|---|---|---|---|---|---|---|---|---|---|---|---|---|---|---|---|---|
| | Unadjusted | | | | Adjusted | | | | Unadjusted | | | | Adjusted | | | |
| | B (SE) | β | p | BH | B (SE) | β | p | BH | B (SE) | β | p | BH | B (SE) | β | p | BH |
| **GM-CSF** | -0.391 (0.129) | -0.173 | 0.003 | 0.024 | -0.401 (0.142) | -0.178 | 0.005 | 0.044 | -0.180 (0.099) | -0.103 | 0.070 | 0.253 | -0.202 (0.106) | -0.115 | 0.057 | 0.205 |
| **IFN-γ** | -0.043 (0.116) | -0.021 | 0.714 | 0.890 | -0.072 (0.127) | -0.039 | 0.572 | 0.822 | 0.077 (0.074) | 0.059 | 0.298 | 0.597 | 0.048 (0.079) | 0.032 | 0.546 | 0.702 |
| **IL-10** | -0.207 (0.096) | -0.124 | 0.032 | 0.114 | -0.215 (0.105) | -0.13 | 0.041 | 0.123 | -0.222 (0.076) | -0.163 | 0.004 | 0.071 | -0.257 (0.081) | -0.191 | 0.001 | 0.018 |
| **IL-12p70** | -0.280 (0.110) | -0.145 | 0.012 | 0.071 | -0.270 (0.121) | -0.141 | 0.025 | 0.090 | -0.138 (0.070) | -0.112 | 0.049 | 0.253 | -0.152 (0.074) | -0.112 | 0.039 | 0.176 |
| **IL-13** | -0.048 (0.180) | -0.015 | 0.791 | 0.890 | -0.027 (0.197) | -0.009 | 0.891 | 0.891 | -0.033 (0.126) | -0.015 | 0.791 | 0.949 | -0.096 (0.132) | -0.048 | 0.468 | 0.648 |
| **IL-1β** | -0.140 (0.109) | -0.056 | 0.333 | 0.666 | -0.140 (0.120) | -0.075 | 0.244 | 0.496 | -0.075 (0.087) | -0.049 | 0.387 | 0.624 | -0.127 (0.092) | -0.082 | 0.166 | 0.427 |
| **IL-2** | -0.266 (0.121) | -0.126 | 0.029 | 0.114 | -0.308 (0.133) | -0.147 | 0.021 | 0.090 | -0.146 (0.079) | -0.105 | 0.064 | 0.253 | -0.174 (0.084) | -0.125 | 0.037 | 0.176 |
| **IL-4** | -0.316 (0.154) | -0.118 | 0.041 | 0.123 | -0.398 (0.169) | -0.148 | 0.019 | 0.090 | -0.006 (0.119) | -0.003 | 0.957 | 0.976 | -0.058 (0.127) | -0.03 | 0.645 | 0.774 |
| **IL-5** | 0.145 (0.112) | 0.075 | 0.194 | 0.437 | 0.142 (0.123) | 0.072 | 0.248 | 0.496 | 0.003 (0.086) | 0.002 | 0.970 | 0.976 | -0.066 (0.090) | -0.054 | 0.465 | 0.648 |
| **IL-6** | -0.042 (0.155) | -0.016 | 0.787 | 0.890 | -0.080 (0.170) | -0.031 | 0.639 | 0.822 | 0.154 (0.116) | 0.076 | 0.184 | 0.413 | 0.117 (0.122) | 0.052 | 0.338 | 0.608 |
| **IL-7** | -0.077 (0.089) | -0.050 | 0.387 | 0.696 | -0.099 (0.097) | -0.066 | 0.308 | 0.554 | -0.050 (0.061) | -0.046 | 0.416 | 0.624 | -0.065 (0.064) | -0.062 | 0.314 | 0.608 |
| **IL-8** | -0.083 (0.120) | 0.040 | 0.492 | 0.737 | -0.095 (0.133) | -0.045 | 0.474 | 0.776 | 0.089 (0.128) | 0.039 | 0.488 | 0.676 | 0.027 (0.136) | 0.01 | 0.842 | 0.892 |
| **TNF-α** | -0.001 (0.081) | -0.001 | 0.992 | 0.992 | 0.032 (0.089) | 0.022 | 0.721 | 0.824 | 0.002 (0.080) | 0.002 | 0.976 | 0.976 | -0.005 (0.085) | -0.001 | 0.952 | 0.952 |
| **NGAL** | -0.118 (0.060) | -0.113 | 0.051 | 0.131 | -0.099 (0.066) | -0.095 | 0.135 | 0.347 | -0.088 (0.065) | -0.077 | 0.175 | 0.413 | -0.070 (0.068) | -0.062 | 0.305 | 0.608 |
| **MMP-9** | 0.035 (0.083) | 0.025 | 0.670 | 0.890 | 0.031 (0.090) | 0.018 | 0.733 | 0.824 | 0.033 (0.071) | 0.027 | 0.641 | 0.824 | 0.019 (0.075) | 0.014 | 0.804 | 0.892 |
| **YKL40** | -0.041 (0.056) | -0.042 | 0.465 | 0.737 | 0.013 (0.060) | 0.016 | 0.823 | 0.871 | 0.128 (0.074) | 0.097 | 0.087 | 0.262 | 0.174 (0.079) | 0.133 | 0.027 | 0.176 |
| **sCD163** | 0.008 (0.059) | 0.008 | 0.893 | 0.945 | 0.031 (0.065) | 0.032 | 0.638 | 0.822 | 0.103 (0.055) | 0.106 | 0.061 | 0.253 | 0.098 (0.058) | 0.1 | 0.091 | 0.273 |
| **sCD14** | 0.203 (0.037) | 0.300 | 0.000 | 0.000 | 0.225 (0.041) | 0.333 | <.001 | 0.018 | 0.032 (0.038) | 0.048 | 0.401 | 0.624 | 0.030 (0.040) | 0.045 | 0.455 | 0.648 |
| | **3 years** | | | | | | | | **5 years** | | | | | | | |
| | **Unadjusted** | | | | **Adjusted** | | | | **Unadjusted** | | | | **Adjusted** | | | |
| | B (SE) | β | p | BH | B (SE) | β | p | BH | B (SE) | β | p | BH | B (SE) | β | p | BH |
| **GM-CSF** | -0.138 (0.108) | -0.081 | 0.200 | 0.761 | -0.231 (0.117) | -0.136 | 0.048 | 0.288 | -0.030 (0.105) | -0.017 | 0.776 | 0.996 | -0.086 (0.113) | -0.051 | 0.446 | 0.825 |
| **IFN-γ** | -0.024 (0.082) | -0.019 | 0.769 | 0.883 | -0.025 (0.090) | -0.018 | 0.782 | 0.938 | -0.021 (0.082) | -0.016 | 0.799 | 0.996 | -0.051 (0.088) | -0.035 | 0.565 | 0.844 |
| **IL-10** | -0.180 (0.066) | -0.170 | 0.007 | 0.121 | -0.235 (0.072) | -0.224 | 0.001 | 0.018 | -0.205 (0.087) | -0.142 | 0.020 | 0.163 | -0.258 (0.094) | -0.176 | 0.006 | 0.111 |

| | | | | | | | | | | | | | | | | |
|---|---|---|---|---|---|---|---|---|---|---|---|---|---|---|---|---|
| **IL-12p70** | 0.038 (0.070) | 0.034 | 0.588 | 0.842 | -0.004 (0.076) | 0.005 | 0.961 | 0.961 | 0.000 (0.067) | 0.000 | 0.996 | 0.996 | -0.043 (0.072) | -0.036 | 0.546 | 0.844 |
| **IL-13** | -0.152 (0.189) | -0.051 | 0.423 | 0.761 | -0.173 (0206) | -0.058 | 0.401 | 0.722 | -0.227 (0.187) | -0.074 | 0.225 | 0.676 | -0.229 (0.201) | -0.076 | 0.254 | 0.763 |
| **IL-1β** | -0.069 (0.067) | -0.065 | 0.303 | 0.761 | -0.124 (0.073) | -0.116 | 0.883 | 0.401 | -0.012 (0.075) | -0.009 | 0.877 | 0.996 | -0.021 (0.081) | -0.017 | 0.797 | 0.844 |
| **IL-2** | 0.033 (0.074) | 0.028 | 0.660 | 0.849 | -0.012 (0.080) | -0.008 | 0.883 | 0.961 | -0.036 (0.087) | -0.025 | 0.681 | 0.996 | -0.096 (0.092) | -0.067 | 0.299 | 0.769 |
| **IL-4** | -0.177 (0.163) | -0.068 | 0.279 | 0.761 | -0.196 (0.176) | -0.076 | 0.267 | 0.722 | -0.245 (0.169) | -0.088 | 0.148 | 0.533 | -0.239 (0.181) | -0.088 | 0.187 | 0.674 |
| **IL-5** | -0.224 (0.093) | -0.149 | 0.017 | 0.157 | -0.267 (0.102) | -0.182 | 0.009 | 0.081 | -0.227 (0.102) | -0.134 | 0.027 | 0.163 | -0.220 (0.110) | -0.133 | 0.046 | 0.277 |
| **IL-6** | -0.138 (0.108) | -0.081 | 0.200 | 0.761 | -0.231 (0.117) | -0.136 | 0.048 | 0.722 | -0.030 (0.105) | -0.017 | 0.776 | 0.401 | -0.086 (0.113) | -0.051 | 0.446 | 0.674 |
| **IL-7** | -0.041 (0.068) | -0.038 | 0.546 | 0.842 | -0.064 (0.073) | -0.059 | 0.383 | 0.722 | -0.009 (0.079) | -0.007 | 0.913 | 0.996 | -0.041 (0.085) | -0.032 | 0.627 | 0.844 |
| **IL-8** | -0.141 (0.169) | -0.052 | 0.407 | 0.761 | -0.100 (0.184) | -0.037 | 0.587 | 0.852 | 0.080 (0.175) | 0.028 | 0.646 | 0.996 | -0.071 (0.187) | -0.027 | 0.705 | 0.844 |
| **TNF-α** | -0.016 (0.071) | -0.014 | 0.820 | 0.883 | 0.033 (0.077) | 0.031 | 0.663 | 0.852 | 0.060 (0.089) | 0.041 | 0.503 | 0.996 | -0.031 (0.095) | -0.022 | 0.744 | 0.844 |
| **NGAL** | -0.030 (0.059) | -0.032 | 0.608 | 0.842 | 0.006 (0.064) | 0.008 | 0.929 | 0.961 | 0.026 (0.052) | -0.031 | 0.614 | 0.996 | -0.047 (0.056) | -0.55 | 0.406 | 0.825 |
| **MMP-9** | -0.001 (0.089) | 0.001 | 0.992 | 0.992 | 0.046 (0.098) | 0.032 | 0.637 | 0.852 | 0.208 (0.091) | -0.138 | 0.023 | 0.163 | 0.241 (0.097) | 0.16 | 0.013 | 0.118 |
| **YKL40** | -0.017 (0.082) | -0.013 | 0.834 | 0.883 | 0.060 (0.089) | 0.049 | 0.502 | 0.821 | -0.005 (0.080) | -0.004 | 0.954 | 0.996 | 0.064 (0.086) | 0.049 | 0.458 | 0.825 |
| **sCD163** | -0.057 (0.063) | -0.057 | 0.370 | 0.761 | -0.074 (0.068) | -0.072 | 0.281 | 0.722 | 0.035 (0.059) | 0.036 | 0.557 | 0.996 | 0.017 (0.064) | 0.019 | 0.793 | 0.844 |
| **sCD14** | 0.072 (0.044) | 0.102 | 0.105 | 0.633 | 0.071 (0.041) | 0.333 | <.001 | 0.511 | -0.009 (0.041) | -0.013 | 0.826 | 0.996 | -0.003 (0.045) | 0.005 | 0.944 | 0.944 |

Adjusted for age, sex, preterm, socioeconomic status, alcohol use, maternal age, maternal body mass index (BMI), maternal HIV. Abbreviations: Benjamini-Hochberg corrected p-value, BH; Granulocyte-macrophage colony-stimulating factor, GM-CSF; interferon-γ, IFN-γ; interleukin, IL; tumor necrosis factor-α, TNF-α; neutrophil gelatinase-associated lipocalin, NGAL; metalloproteinase-9, MMP-9; soluble cluster of differentiation, sCD; chitinase-3-like protein 1, YKL-40.

**Supplementary Table 5.** Relationship between maternal ART initiation and cytokine levels in children

| | 6 weeks | | | | | | 2 years | | | | | |
|---|---|---|---|---|---|---|---|---|---|---|---|---|
| | | **Pre-pregnancy** | | **During Pregnancy** | | | | **Pre-pregnancy** | | **During Pregnancy** | | |
| | t | lower | upper | lower | upper | p | t | lower | upper | lower | upper | p |
| **GM-CSF** | -1.219 | 2.925 | 20.198 | 5.468 | 17.998 | 0.225 | 1.241 | 62.79875 | 198.58750 | 39.74375 | 163.51125 | 0.217 |
| **IFN-γ** | -1.314 | 1.528 | 10.979 | 2.795 | 13.098 | 0.191 | 0.080 | 6.74250 | 16.19750 | 5.42250 | 17.43250 | 0.936 |
| **IL-10** | -0.776 | 5.50375 | 20.72750 | 7.24625 | 18.70875 | 0.439 | 0.968 | 12.74875 | 25.35375 | 9.59500 | 22.08500 | 0.335 |
| **IL-12p70** | -1.692 | 0.51250 | 3.39125 | 0.92250 | 3.13000 | 0.093 | 0.165 | 2.85125 | 5.62125 | 2.95000 | 5.38500 | 0.869 |
| **IL-13** | -0.465 | 0.88250 | 7.65000 | 1.05500 | 6.79250 | 0.643 | -0.781 | 4.28625 | 20.78750 | 5.07500 | 22.09250 | 0.436 |
| **IL-1β** | -0.227 | 0.42250 | 1.62875 | 0.37250 | 1.55250 | 0.820 | 0.036 | 1.09750 | 2.66000 | 1.19125 | 2.92375 | 0.971 |
| **IL-2** | -0.881 | 0.29774 | 2.03000 | 0.49500 | 1.92250 | 0.380 | 0.339 | 1.63125 | 3.75250 | 1.33375 | 3.57750 | 0.735 |
| **IL-4** | -0.215 | 3.47375 | 29.39625 | 4.55750 | 23.12250 | 0.830 | 0.198 | 18.87875 | 54.41625 | 11.94625 | 60.95375 | 0.843 |
| **IL-5** | -1.297 | 0.67239 | 2.57625 | 1.30000 | 2.65250 | 0.197 | -0.102 | 2.40375 | 5.75125 | 2.64375 | 5.67875 | 0.919 |
| **IL-6** | -0.827 | 0.53059 | 4.73250 | 0.66500 | 3.85000 | 0.410 | -0.653 | 1.98750 | 6.33375 | 2.01500 | 8.25500 | 0.515 |
| **IL-7** | -1.276 | 3.02250 | 8.84250 | 3.98750 | 9.89500 | 0.204 | -0.579 | 6.43125 | 13.73500 | 6.82625 | 14.58625 | 0.564 |
| **IL-8** | 0.054 | 5.05500 | 14.61875 | 4.34000 | 17.21500 | 0.957 | 0.172 | 7.19000 | 26.54500 | 6.12000 | 34.17000 | 0.864 |
| **TNF-α** | -0.154 | 14.54375 | 29.75125 | 13.49875 | 35.89250 | 0.878 | 0.328 | 9.52125 | 18.91125 | 7.51125 | 18.24000 | 0.743 |
| **NGAL** | 0.059 | 60.19250 | 136.75844 | 63.21692 | 120.97922 | 0.953 | -0.814 | 100.43115 | 200.11970 | 107.38150 | 224.57924 | 0.417 |
| **MMP-9** | 1.663 | 301.68716 | 812.86372 | 279.94600 | 658.19481 | 0.099 | -0.935 | 602.36496 | 1238.68487 | 492.09355 | 1512.96369 | 0.352 |
| **YKL40** | 0.227 | 23.50544 | 39.52352 | 22.48642 | 42.41744 | 0.821 | -1.247 | 19.12673 | 39.00854 | 19.78146 | 57.87361 | 0.215 |
| **sCD163** | -0.625 | 374.66726 | 858.85223 | 411.61744 | 846.68137 | 0.533 | -0.629 | 507.58261 | 1008.77589 | 501.75535 | 978.10922 | 0.531 |
| **sCD14** | -1.205 | 1411.06228 | 2116.94154 | 1515.01049 | 2266.26428 | 0.230 | 0.088 | 1704.10141 | 2815.53142 | 1586.20879 | 2681.02934 | 0.930 |
| | **3 years** | | | | | | **5 years** | | | | | |
| | | **Pre-pregnancy** | | **During Pregnancy** | | | | **Pre-pregnancy** | | **During Pregnancy** | | |
| | **t** | **lower** | **upper** | **lower** | **upper** | **p** | **t** | **lower** | **upper** | **lower** | **upper** | **p** |
| **GM-CSF** | -0.229 | 32.80250 | 131.69875 | 38.83000 | 132.47875 | 0.819 | 0.837 | 38.922500 | 131.611250 | 35.060000 | 118.062500 | 0.405 |
| **IFN-γ** | -0.092 | 14.12875 | 31.06625 | 13.78375 | 30.65125 | 0.927 | 0.324 | 12.71250 | 24.96750 | 12.10000 | 23.83750 | 0.747 |
| **IL-10** | -0.704 | 6.75875 | 13.00500 | 7.72125 | 13.03500 | 0.483 | 0.736 | 4.79250 | 11.63500 | 3.99500 | 9.28500 | 0.464 |
| **IL-12p70** | 0.242 | 2.62625 | 5.07625 | 2.84750 | 5.36125 | 0.810 | 0.296 | 2.16500 | 4.62250 | 2.33500 | 4.25250 | 0.768 |
| **IL-13** | -0.087 | 3.65250 | 24.84000 | 4.19875 | 20.56875 | 0.931 | 0.394 | 1.96875 | 35.14875 | 2.31000 | 23.31750 | 0.695 |

| | | | | | | | | | | | | |
|---|---|---|---|---|---|---|---|---|---|---|---|---|
| **IL-1β** | -0.406 | 1.65625 | 3.62250 | 2.04250 | 3.39250 | 0.686 | 0.829 | 1.86875 | 3.71000 | 1.83000 | 3.73500 | 0.409 |
| **IL-2** | 0.216 | 1.66500 | 3.95875 | 1.77750 | 3.39375 | 0.830 | 0.815 | 1.53750 | 3.86250 | 1.37750 | 3.31750 | 0.417 |
| **IL-4** | 0.106 | 15.29250 | 111.38250 | 20.45875 | 95.82625 | 0.916 | 0.190 | 11.43875 | 103.46500 | 12.94500 | 59.98000 | 0.850 |
| **IL-5** | 0.475 | 2.03625 | 4.18375 | 1.97125 | 4.30750 | 0.636 | 0.511 | 2.01250 | 4.15000 | 1.38750 | 4.19500 | 0.611 |
| **IL-6** | 0.838 | 1.46000 | 9.44375 | 1.47125 | 7.76375 | 0.404 | 0.547 | 0.77625 | 7.67750 | 0.93000 | 5.25750 | 0.586 |
| **IL-7** | -0.428 | 6.72625 | 14.09500 | 7.27875 | 14.57250 | 0.670 | -0.008 | 6.09375 | 14.56750 | 6.47500 | 13.95750 | 0.994 |
| **IL-8** | 0.522 | 8.07250 | 31.26250 | 5.73125 | 31.53250 | 0.603 | -0.140 | 5.75375 | 49.44125 | 6.97000 | 33.89250 | 0.889 |
| **TNF-α** | 1.443 | 8.512500 | 16.795000 | 7.916250 | 15.038750 | 0.152 | -0.624 | 8.11375 | 12.33000 | 6.32000 | 14.41500 | 0.534 |
| **NGAL** | -0.532 | 78.91071 | 148.28103 | 85.90210 | 157.69884 | 0.596 | -0.600 | 87.15915 | 138.98447 | 78.47684 | 167.56528 | 0.550 |
| **MMP-9** | -0.420 | 520.28402 | 1471.51176 | 558.51952 | 1740.17407 | 0.675 | 0.206 | 610.22076 | 2158.08923 | 623.92661 | 1619.22117 | 0.837 |
| **YKL40** | -1.532 | 13.25369 | 41.30051 | 17.86009 | 43.47144 | 0.129 | -0.802 | 14.46031 | 26.90438 | 14.23816 | 36.48530 | 0.424 |
| **sCD163** | 0.204 | 413.76483 | 1029.18006 | 427.09382 | 827.93998 | 0.839 | -0.935 | 420.03691 | 711.10073 | 404.74781 | 758.01526 | 0.352 |
| **sCD14** | -1.145 | 1774.96485 | 2843.44916 | 1985.62967 | 2880.50353 | 0.255 | 0.651 | 1735.50079 | 2541.69182 | 1734.26060 | 2473.33699 | 0.516 |

Adjusted for age, sex, preterm, socioeconomic status, alcohol use, maternal age, maternal body mass index (BMI), maternal HIV. Abbreviations: Granulocyte-macrophage colony-stimulating factor, GM-CSF; interferon-γ, IFN-γ; interleukin, IL; tumor necrosis factor-α, TNF-α; neutrophil gelatinase-associated lipocalin, NGAL; metalloproteinase-9, MMP-9; soluble cluster of differentiation, sCD; chitinase-3-like protein 1, YKL-40.
